# Interaction of septin 7 and DOCK8 in equine lymphocytes reveals novel insights into signaling pathways associated with autoimmunity

**DOI:** 10.1101/251819

**Authors:** Melanie Schauer, Kristina J H Kleinwort, Roxane L Degroote, Carmen Wiedemann, Elisabeth Kremmer, Stefanie M Hauck, Cornelia A Deeg

**Affiliations:** Chair for Animal Physiology, Department of Veterinary Sciences, LMU Munich, 80539 Munich, Germany; Institute for Molecular Immunology, Helmholtz Center Munich, German Research Center for Environmental Health GmbH, 81377 Munich, Germany; Research Unit Protein Science, Helmholtz Center Munich, German Research Center for Environmental Health GmbH, 80939 Munich, Germany

## Abstract

The GTP-binding protein septin 7 is involved in various cellular processes, including cytoskeleton organization, migration and the regulation of cell shape. Septin 7 function in lymphocytes, however, is poorly characterized. Since the intracellular signaling role of septin 7 is dependent on its interaction network, interaction proteomics was applied to attain novel knowledge about septin 7 function in hematopoietic cells. Our previous finding of decreased septin 7 expression in blood-derived lymphocytes in ERU, a spontaneous animal model for autoimmune uveitis in man, extended the role of septin 7 to a potential key player in autoimmunity. Here, we revealed novel insights into septin 7 function by identification of DOCK8 as an interaction partner in primary blood-derived lymphocytes. Since DOCK8 is associated with important immune functions, our finding of significantly decreased DOCK8 expression and altered DOCK8 interaction network in ERU might explain changes in immune response and shows the contribution of DOCK8 in pathomechanisms of spontaneous autoimmune diseases. Moreover, our analyses revealed insights in DOCK8 function, by identifying the signal transducer ILK as a DOCK8 interactor in lymphocytes. Our finding of the enhanced enrichment of ILK in ERU cases indicates a deviant influence of DOCK8 on inter- and intracellular signaling in autoimmune disease.

## INTRODUCTION

Septins are GTP-binding cytoskeletal proteins involved in various cellular processes, including cytokinesis, membrane dynamics and regulation of cell shape^1,2^ They are highly conserved from yeast to humans. One unique feature of septins is the ability to self-assemble and form filaments that serve as scaffolds for protein recruitment^3^. The assembly of septin filaments consists of hexamers and octamers which are formed by septin subunits^4^, all including the essential group member septin 7 ^5,6^. Due to its pivotal role in the formation of septin filaments, the depletion of septin 7 leads to destabilization of the core septin cytoskeleton in mammalian cells^1^.

We previously found septin 7 protein downregulated in CD4^+^ T cells of equine recurrent uveitis (ERU) cases^7^ ERU represents the only spontaneous model for autoimmune uveitis in humans^8^ and is characterized by remitting painful attacks of the inner eye, caused by infiltration of immune cells^9^, eventually leading to vision loss. During uveitic attacks, autoreactive CD4^+^ T-lymphocytes cross the blood-retina-barrier, infiltrating ciliary body, choroid and retina and destruct retinal tissue^9,10^. The initiating pathomechanism causing these peripheral lymphocytes to target intraocular tissue is still poorly understood. One possible explanation might be their altered septin 7 expression level leading to changed migratory behavior and deviant activation of T cells^7^ In murine primary T cells, knock-down of septin 7 led to enhanced blebbing and protrusion formation, coming from a loss of membrane stabilization^2^. Moreover, T cells lacking septin 7 showed a significantly higher transmigration rate through very small pores^11^. This observation might result from relaxation of the cell cortex, allowing efficient migration through especially narrow pores^11^. In autoimmune uveitis migration of T cells over blood-retina-barrier into the eye is a key driver of pathology^12–14^ Since decreased septin 7 expression might result in the ability of T cells to cross this barrier, septin 7 might take a pivotal role in cellular processes that regulate immune functions such as cell signaling or cell migration.

However, the impact of expression changes or functional deviations of septin 7 in lymphocytes in course of autoimmune disease was not described to date. Since interaction networks of septins are functionally crucial due to their signal-mediating role in cellular processes^15^, unraveling the septin 7 interactome in lymphocytes could contribute to a better understanding of septin 7 function. Therefore, we investigated septin 7 protein networks in primary blood-derived lymphocytes (PBL) with a special focus on immune response related candidates and their functions in autoimmune reactions.

## RESULTS

### Detection of DOCK8 as a novel interaction partner of septin 7

Since the impact of changes in septin 7 protein levels in lymphocytes is yet to be unraveled, we further characterized septin 7 function by identifying septin 7-interacting partners in PBL of healthy cases with immune precipitation, followed by label-free LC-MS/MS mass spectrometry. A total of 20 septin 7 interacting proteins could be detected in this experiment (Table 1). Amongst these, known septin 7 interactors from studies in other models were identified, e.g. septin 2, septin 6 and the small GTPase CDc42^16,17^ Additionally, we detected 10 novel septin 7 interactors in PBL, not described before to our knowledge (Table 1).

**Table 1:** Interacting proteins of septin 7 in PBL of healthy controls. List of septin 7 interactors with enrichment factor ≥ 18 detected by co-immunoprecipitation and identified by LC-MS/MS mass spectrometry. Column 1 (Protein name) shows name of protein. Column 2 (Gene name) shows gene name of identified protein. Column 3 (Accession) shows matching protein accession number, as listed in Ensembl Horse protein database (http://www.ensembl.org/Equus_caballus/Info/Index). Column 4 (Enrichment factor) shows the ratio of normalized protein abundance from enrichment compared to respective isotype controls.

| Protein name | Gene name | Accession | Enrichment factor |
| --- | --- | --- | --- |
| Actin related protein 2/3 complex, subunit 2, 34kDa | ARPC2 | ENSECAP00000009522 | **3464,2** |
| Histone H2A | LOC100053378 | ENSECAP00000001117 | **101,3** |
| Cell division cycle 42 | CDC42 | ENSECAP00000003324 | **76,6** |
| Capping protein (actin filament) muscle Z-line, alpha 1 | CAPZA1 | ENSECAP00000014750 | **71,3** |
| Myosin, light chain 6, alkali, smooth muscle and non-muscle | MYL6 | ENSECAP00000016002 | **69,2** |
| Actin, cytoplasmic 2 | ACTB | ENSECAP00000016124 | **51,8** |
| Septin 6 | SEP6 | ENSECAP00000021088 | **48,5** |
| Histone cluster 1, H2bj | HIST1H2BJ | ENSECAP00000001004 | **46,8** |
| Vimentin | VIM | ENSECAP00000004638 | **41,4** |
| Fibrinogen alpha chain | FGA | ENSECAP00000008007 | **40,8** |
| C-reactive protein, pentraxin-related | CRP | ENSECAP00000022415 | **40,7** |
| Actinin, alpha 1 | ACTN1 | ENSECAP00000017987 | **40,3** |
| Septin 7 | SEP7 | ENSECAP00000020433 | **36,0** |
| Dedicator of cytokinesis 8 | DOCK8 | ENSECAP00000010583 | **32,2** |
| Myosin IG | MYO1G | ENSECAP00000016084 | **28,2** |
| Flotillin 2 | FLOT2 | ENSECAP00000010141 | **23,5** |
| Protein S100-A8 | S100A8 | ENSECAP00000007199 | **22,7** |
| Tropomodulin 3 (ubiquitous) | TMOD3 | ENSECAP00000018986 | **20,8** |
| Cathepsin G | CTSG | ENSECAP00000003203 | **20,4** |
| Septin 2 | SEP2 | ENSECAP00000015383 | **18,8** |

One of these interaction partners, dedicator of cytokinesis 8 (DOCK8), regulates various immune functions in lymphocytes^18–20^. DOCK8 was shown to be involved in immune synapse formation, lymphocyte migration and organization of cell shape^19^ We confirmed DOCK8 in septin 7-immunoprecipitates (Figure 1B)^21^. In the reverse experiment, immunoprecipitation of DOCK8 also clearly co-precipitated septin 7. DOCK8 as well as septin 7 were not detected in immunoprecipitates of the negative control MALT1^21^ (Figure 1C). These results point to a protein-protein-interaction of septin 7 and DOCK8.

**Fig. 1:**
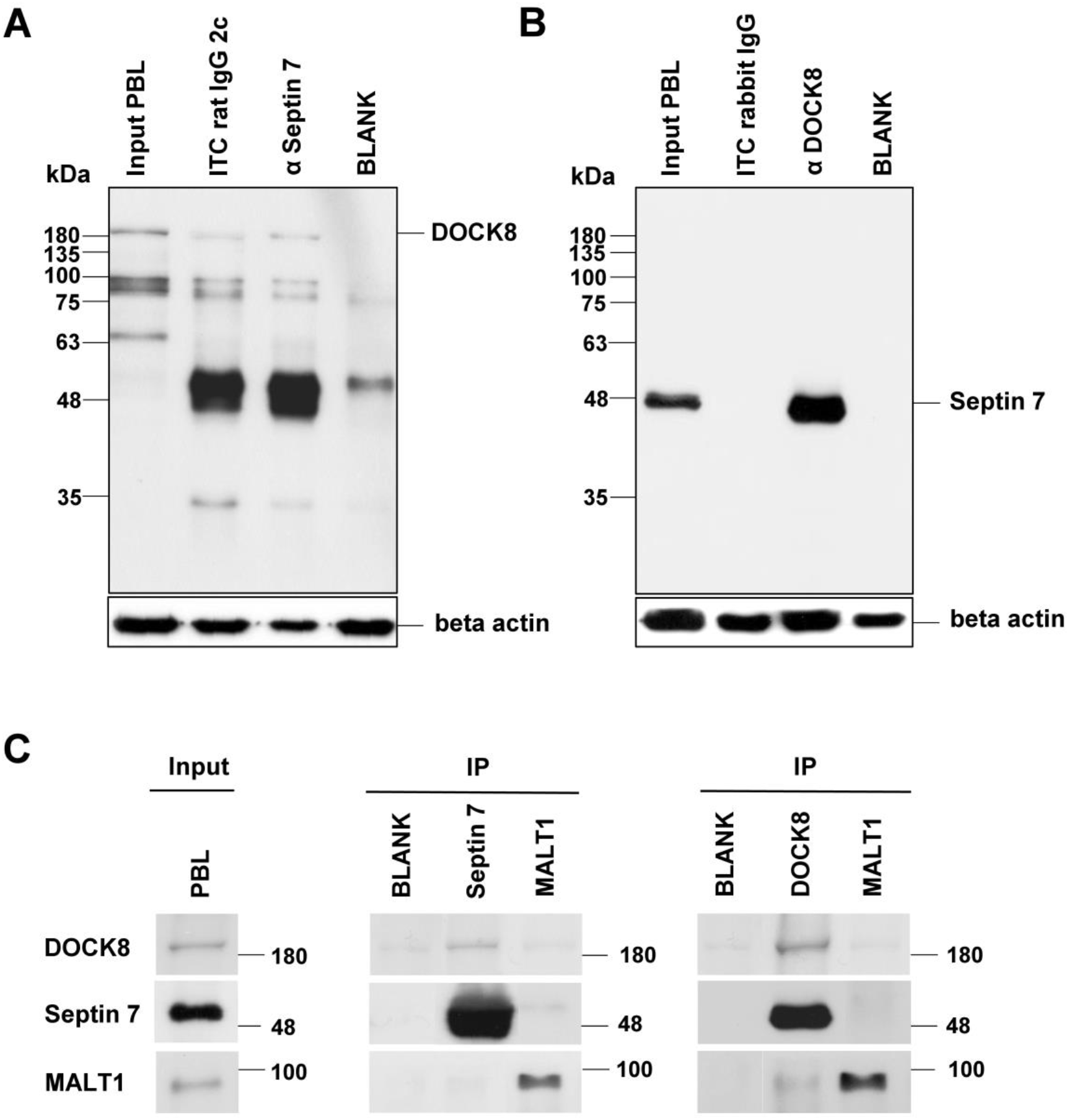
Septin 7 interacted with DOCK8 in equine PBL. Representative western blots of immunoprecipitations of septin 7 (A) and DOCK8 (B). DOCK8 was detected at 190 kDA in input lysates of PBL (input PBL) as well as after immunoprecipitation of septin 7 (A). Septin 7 was detected at 49 kDA in input lysates of PBL (input PBL) as well as after immunoprecipitation of DOCK8, verifying an interaction between these two proteins. In respective isotype controls (rabbit IgG; rat IgG 2c), blank (IP buffer) and negative control MALT1; neither septin 7 nor DOCK8 was detected (C).

### DOCK8 was highly expressed in all lymphocyte subsets

To gain further insights in physiological DOCK8 expression pattern, we analyzed DOCK8 expression levels in CD4^+^ T cells, CD8^+^ T cells and in B cells of horses using flow cytometry. DOCK8 was expressed in all lymphocyte subsets with high intensity (Figure 2), which points to similarities across species, because this was also shown in mice and man^18^. The protein level of DOCK8 in B cells was significantly higher as compared to T cells (two-fold; Figure 2). This DOCK8 expression pattern was never described in other species, since proteomic data generated in human lymphocytes showed equal expression pattern of DOCK8 in T cells and B cells^22^

**Fig. 2:**
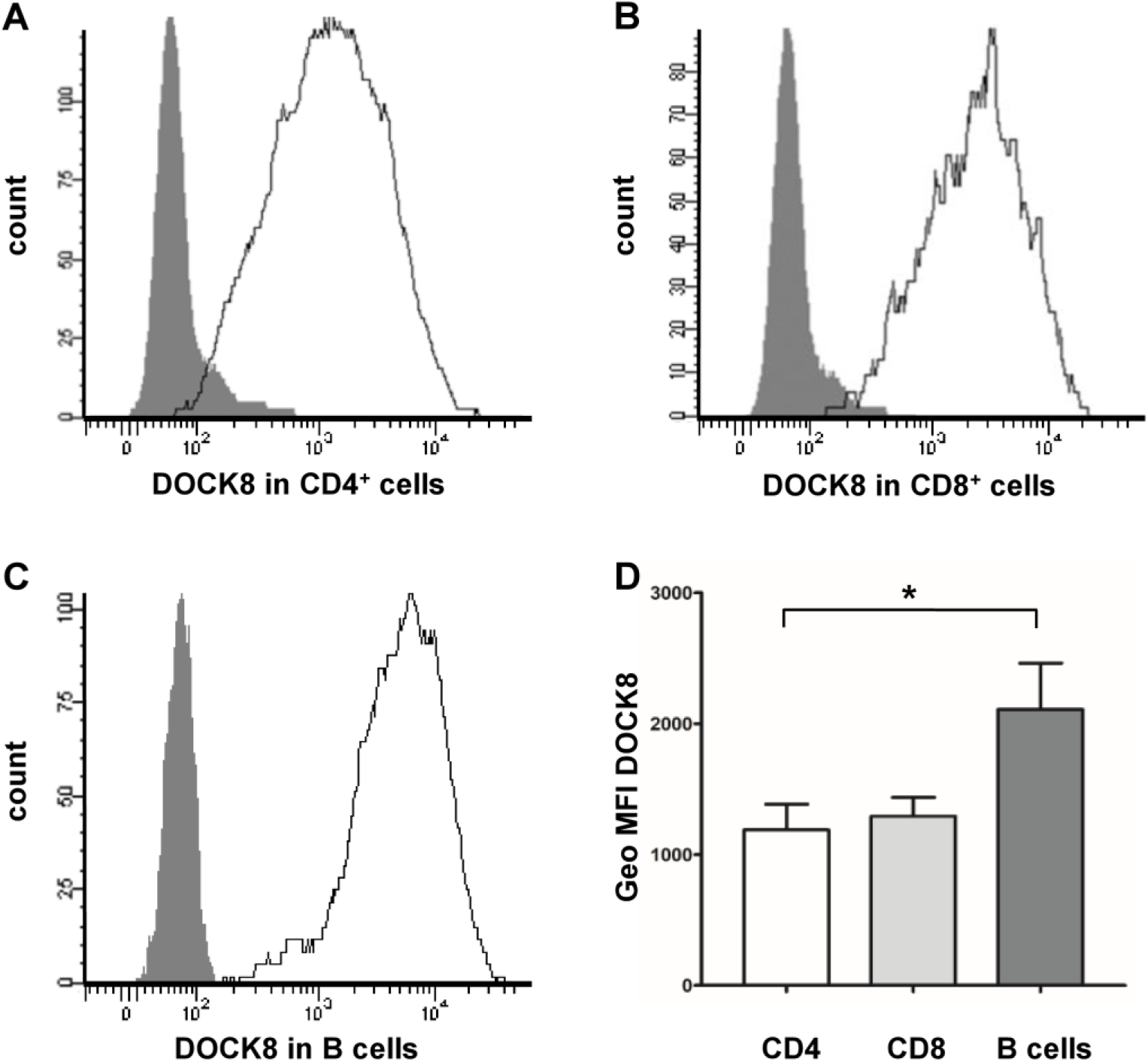
Characterization of DOCK8 expression in horse lymphocytes. (A) Representative DOCK8 expression in different lymphocyte subsets. Flow cytometry analyses demonstrate that DOCK8 is expressed in CD4^+^ T cells, CD8+ T cells and in B cells (white histograms) with high intensity. White histograms represent specific stainings, grey histograms represent respective isotype controls. Y-axis show the amount of cells included in measurement; x-axis show the intensity of DOCK8 staining. (B) Statistical analyses of geo mean fluorescence intensity of DOCK8 expression in lymphocyte subsets of 12 healthy controls. While there was no difference between DOCK8 expression intensity in CD4^+^ T cells (white column) and CD8+ T cells (light grey column), DOCK8 was significantly higher (* = p ≤ 0.05) expressed in B cells (dark grey column) compared to T cells.

### DOCK8 expression in lymphocytes was decreased in autoimmune cases

Next, we analyzed DOCK8 protein levels in lymphocytes of autoimmune cases to detect possible differences in expression. A highly significant (*** = p ≤ 0.001) decrease of DOCK8 expression to 63% (± 28.5% standard deviation (SD)) in ERU compared to healthy controls (100% ± 38.3% SD) was detected by western blot (Figure 3A+3B). To exclude possible differences in cellular composition as a reason for different DOCK8 expression, we controlled respective PBL subset proportions in healthy and ERU cases. There was no significant alteration in cellular compositions detectable between control and ERU group (Figure 3C). Next, we determined if DOCK8 downregulation was predominant in a lymphocyte subset by analyzing DOCK8 expression in CD4^+^ and CD8+ T cells as well as in B cells by flow cytometry. DOCK8 was also present in all lymphocyte subsets of ERU cases, but with considerably decreased levels (geo MFI) in all lymphocyte subsets analyzed (Figure 3D). In T°cells, differences in DOCK8 protein levels were not significant. In B cells, however, the DOCK8 abundance was significantly (* = p ≤ 0.05) decreased to 57% in PBL of ERU cases, compared to a 100% in healthy controls.

**Fig. 3:**
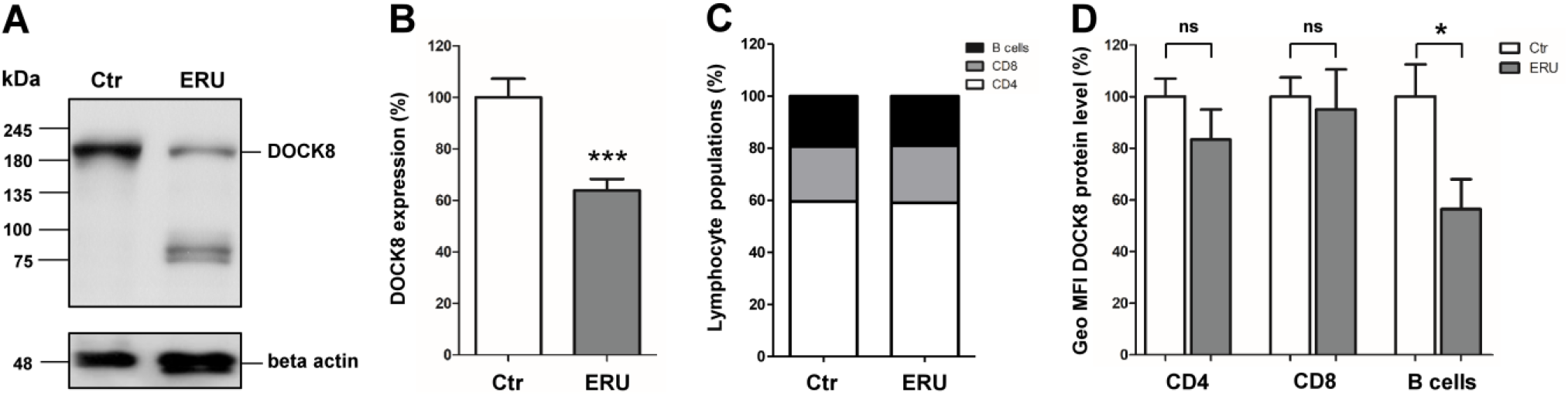
Diminished DOCK8 protein levels in PBL of ERU cases. (A) Representative signal abundances of DOCK8 in PBL of controls (left panel) and ERU (right panel) detected by western blot. DOCK8 signal abundance in PBL of diseased specimen was decreased compared to controls. Lower molecular extra bands result from further reactivities of the polyclonal rabbit anti DOCK8 antibody (B). Compared to healthy controls (n=16), DOCK8 levels in ERU PBL (n=36) were significantly (*** = p ≤ 0.001) decreased to 63% of physiological expression rate (set to a 100%). All protein abundances of DOCK8 were normalized to beta actin. (C) Analyzes of lymphocyte subset composition in healthy controls and autoimmune cases revealed no significant differences. Statistical analysis of DOCK8 expression differences in lymphocyte subsets of healthy controls (white bars; n=13) and ERU cases (grey bars; n=12). DOCK8 expression intensity (geo MFI) was decreased in all lymphocyte subsets of ERU cases. In CD8+ T cells, no considerable DOCK8 expression difference between control PBL and ERU cases could be detected, whereas in CD4^+^ T cells of ERU horses, geo MFI was reduced to 85% compared to healthy controls. In B cells of autoimmune cases, a significant decrease of DOCK8 geo MFI to 57% was detected (* = p ≤ 0.05).

### Unraveling different modes of DOCK8 action in physiological immune response and autoimmunity

Since the functional impact of differential DOCK8 protein levels in autoimmune diseases is unknown, we further investigated potential differences in protein-protein-interaction network of DOCK8 in healthy and autoimmune phenotypes with interaction proteomics. We identified a total of 259 potential DOCK8 interaction partners, that were detectable in all lymphocyte samples analyzed. Interestingly, we could detect remarkable differences in DOCK8-protein enrichment patterns between lymphocytes of healthy and autoimmune case groups. Amongst the 259 interacting proteins, 53 proteins were enriched in both phenotypes, whereas 28 proteins showed enhanced enrichment in lymphocytes of healthy controls and 178 proteins were enriched in lymphocytes of autoimmune cases.

To determine whether DOCK8 functions through different signal transduction pathways in controls and diseased cases, we analyzed DOCK8 interaction network for enrichment of canonical pathways with gene ranker in both phenotypes. Interestingly, we found different signaling pathways with enhanced enrichment in control lymphocytes and in ERU cases (Table 2; Table 3). In PBL of healthy controls, the pathways “ARP2 Actin Related Protein 2 Homolog”, “Small GTP-binding Protein RAC” and “Myosin light chain kinase” were enriched (Table 2). This differed in lymphocytes of ERU cases, where only the pathway “ARP2 Actin Related Protein 2 Homolog” was concurrently enriched (Table 3). Then the two signaling pathways “Integrin” and “Cofilin” were among the top 3 DOCK8-interacting signaling pathways enriched in PBL of autoimmune cases (Table 3). Since proteins comprised in integrin signaling pathway are associated with a wide range of important immune functions, as cell migration or survival, we were especially interested in the role of DOCK8-interactors associated to integrin signaling. This revealed proteins, as talin 1 and integrin-linked kinase (ILK) (Table 3), which we previously found involved in inflammatory processes in ERU cases^23,24^.

**Table 2:** Overrepresented signaling pathways of DOCK8 interacting proteins in PBL of healthy controls. List of the top 3 significantly (p-value) enriched signaling pathways containing DOCK8-interacting proteins enriched in healthy PBL (≥ 2 peptide counts; enrichment factor ≥ 4) analyzed by Genomatix (version 3.7).

| Signaling pathway | *p*-value | No. of proteins observed | List of observed proteins (gene symbols) |
| --- | --- | --- | --- |
| ARP2 Actin Related Protein 2 Homolog (Yeast) | 3,85E-08 | 6 | ACTR2, GSN, VCL, ACTB, ARPC4, ACTR3 |
| Small GTP-binding Protein RAC | 3,13E-04 | 8 | ACTR2, FERMT3, RSU1, DOCK8, CORO1C, IQGAP1, C1QA, PARVB |
| Myosin light chain kinase | 6,35E-04 | 4 | ACTN1, MYO18A, MYH10, MYL9 |

**Table 3:** Overrepresented signaling pathways of DOCK8 interacting proteins in PBL of ERU cases. List of the top 3 significantly (p-value) enriched signaling pathways containing DOCK8-interacting proteins enriched in ERU PBL (≥ 2 peptide counts; enrichment factor ≥ 4) analyzed by Genomatix (version 3.7).

| Signaling pathway | *p*-value | No. of proteins observed | List of observed proteins (gene symbols) |
| --- | --- | --- | --- |
| ARP2 Actin Related Protein 2 Homolog (Yeast) | 8,39E-06 | 6 | VCL, ARPC4, CAPZA1, ACTR3, GSN, ACTB |
| Integrin | 3,87E-04 | 12 | LGALS3BP, ITGB3, VCL, FLNA, FN1, C3, TRF, VWF, ACTN1, TLN1, ILK, C1QA |
| Cofilin | 1,44E-03 | 8 | MYO9B, HSPB1, SEPT7, NEXN, PHB, GSN, SEPT2, ACTB |

### Enhanced expression of ILK, a novel DOCK8 interaction partner in autoimmune cases

Since the role of ILK in autoimmunity is yet to be unraveled, we further characterized the role of ILK in PBL of controls and autoimmune cases. Due to its crucial involvement in signal transduction, chemotaxis and regulation of cell survival^25^, ILK represents a potential key player in ERU. In DOCK8-precipitates, ILK could clearly be detected and in the reverse experiment DOCK8 was also co-precipitated in ILK-precipitate, clearly verifying the protein-protein interaction of both candidates (Figure 4). By comparing ILK protein levels in lymphocytes of healthy and ERU cases, we detected a significant increase of ILK protein expression in PBL of diseased specimen (Figure 5A).

**Fig. 4:**
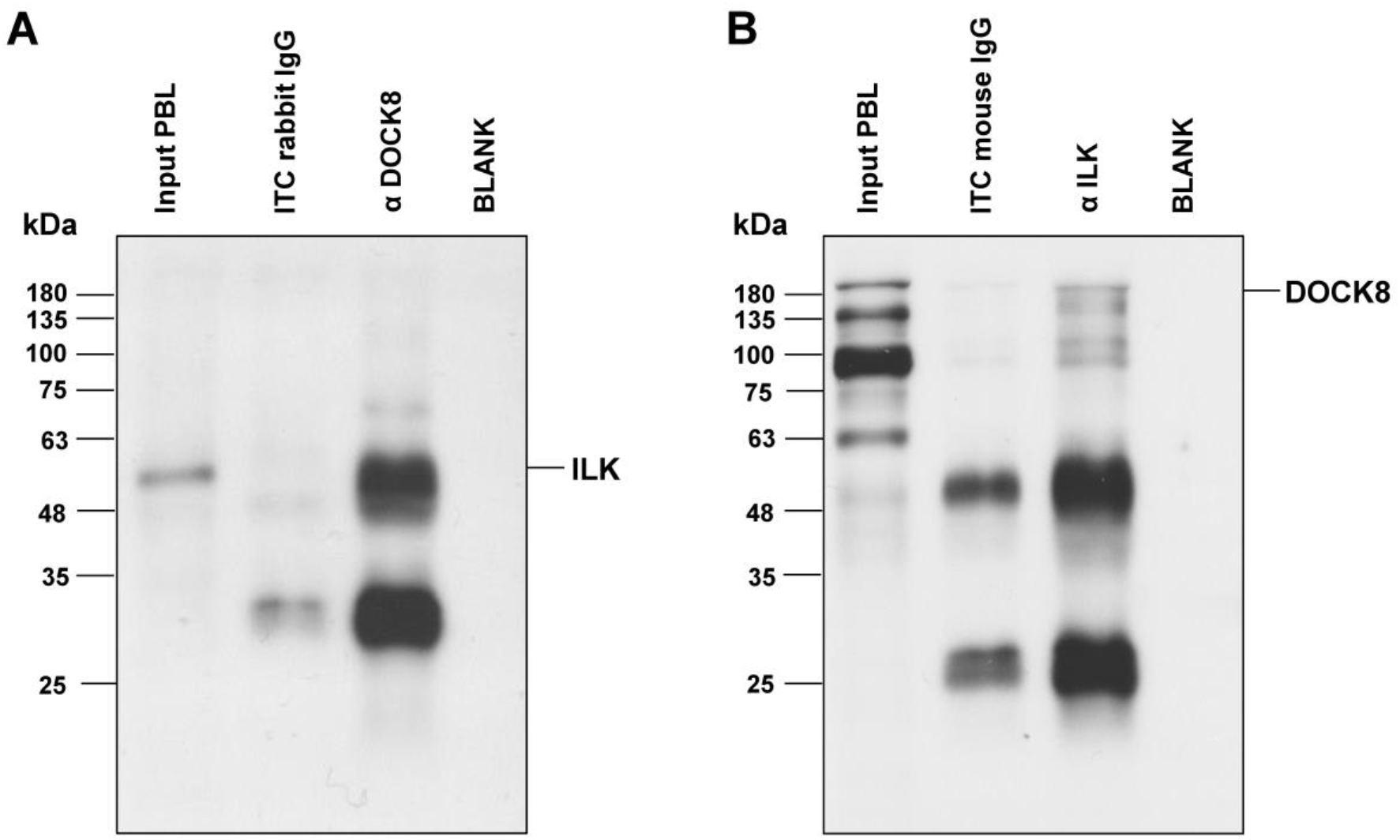
DOCK8 interacted with ILK in equine PBL. Representative western blots of immunoprecipitations of DOCK8 (A) and ILK (B). ILK was detected at 59 kDA in input lysates of PBL (input PBL) as well as after immunoprecipitation of DOCK8 (A). DOCK8 was detected at 190 kDA in input lysates of PBL (input PBL) as well as after immunoprecipitation of ILK, verifying an interaction between these two proteins. In respective isotype controls (rb IgG; mIgG) and blank (IP buffer), neither ILK nor DOCK8 was detected.

**Fig. 5:**
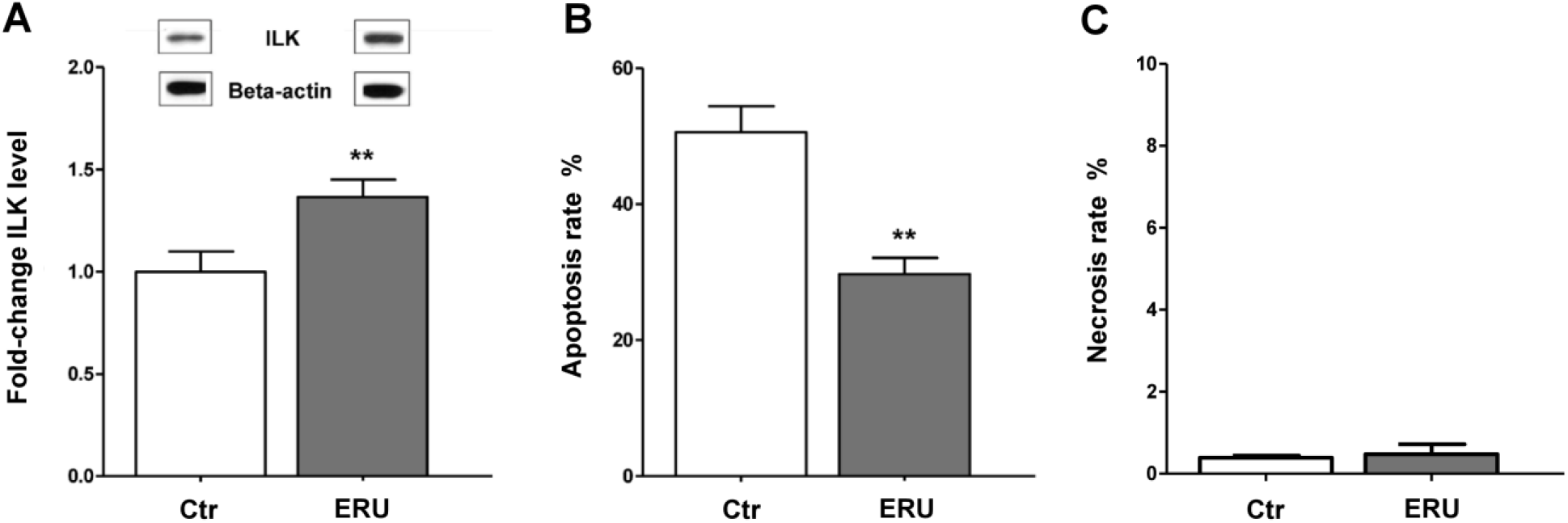
ILK expression in lymphocytes was significantly increased in diseased state and coincidented with increased cell survival. (A) Significantly increased ILK protein levels in ERU cases (n=33) compared to healthy controls (n=18) were verified and quantified by western blots (** = p ≤ 0.01). ILK intensities were normalized to beta actin abundances on respective blots. Representative signal abundances are shown above respective columns. (B) PBL of healthy (biological n=4, technical replicates n=2) and ERU horses (biological replicates n=6), which constitutively differed in ILK expression were cultured at 37°C for 72 hours. Apoptosis rate was determined by measuring the amount of annexin V positive cells in flow cytometry. ERU horses (grey column), expressing ILK significantly higher (1.5-fold), showed a highly significantly reduced amount of apoptotic cells compared to healthy controls (white column) (** = p ≤ 0.01). (C) In contrast, necrosis rate of lymphocytes, determined by the amount of PI positive cells, was not different between healthy controls (white column) and ERU cases (grey bar).

### Lymphocytes with increased ILK expression have higher survival rate

Since ILK was shown to be essential for various cellular functions including the regulation of survival in murine lymphocytes^25^, we hypothesized that increased expression intensity of ILK in spontaneous autoimmune cases might result in a higher survival rate of respective lymphocytes. This could affect immune tolerance and provoke autoaggressive immune reaction. Therefore, we investigated apoptotic rates of lymphocytes of healthy and autoimmune cases, showing a constitutive differential ILK expression level. Increased ILK protein level in lymphocytes of diseased specimen coincided with a significantly (** = p ≤ 0.01) reduced apoptosis rate of these cells (Figure 5B). Whereas PBL of control cases had an average apoptosis rate of 50.6% ± 10.8% SD; apoptosis rate of autoimmune cases was only 29.7% ± 5.8% SD. Necrosis of lymphocytes was equally low in both phenotypes (< 1%) (Figure 5C).

## DISCUSSION

The aim of our study was to extend the knowledge about the role of septin 7 in lymphocytes and in the context of a spontaneous autoimmune disease. We previously observed decreased levels of septin 7 in lymphocytes isolated from ERU cases^7^ Since functional impact of reduced septin 7 levels in lymphocytes and, more generally, in horse cells, was unknown, we chose a protein-protein interaction screening approach to identify proteins potentially relevant for signal transduction pathways triggered by septin 7. We identified a total of 20 septin 7 interacting proteins from primary PBL. Nine of these proteins were previously suggested as septin 7 interactors (Supplementary Table S1), however identified mostly in yeast cells using septin 7 plasmid-driven overexpression^26^. Since our approach based on immunoprecipitation from native cells, the confirmation of previously described interactors suggests similar core units of the septin interactome across cell types and species. Among those reconfirmed septin 7 interactors were proteins shown to be involved in cytokinesis and cell migration in other models, e.g. septin 2 and septin 6^11,26^ (Table 1).

Furthermore, we detected 10 proteins interacting with septin 7 in lymphocytes, which were previously unknown. Amongst those, especially the identification of DOCK8 as a novel septin 7 interaction partner was a promising finding that could help to elucidate pathomechanisms in autoimmune diseases in our opinion. Like septin 7, DOCK8 is involved in the organization of actin cytoskeleton and regulates the structural integrity during immune cell trafficking^19,21^. In particular, DOCK8 guanine nucleotide exchange factor (GEF) activity for CDC42 was shown to be essential for T cell receptor driven actin assembly^21^. Interestingly, CDC42 is a shared interactor of both DOCK8 and septin 7 ^17,27,28^, which we also confirmed by co-immunprecipitation following by mass spectrometry (interaction with septin 7 shown in Table 1, DOCK8-interaction data not shown). This molecule might represent a key driver that mediates the protein-protein-interaction and consequently takes an essential part in cell signaling, involving septin 7 and DOCK8.

Similar to septin 7 ^7^, DOCK8 levels were reduced in lymphocytes of ERU cases (Figure 3) strengthening the functional overlap between these two molecules, which might affect the regulation of immune cell migration. Moreover, investigations on life-threatening DOCK8-defıciency in humans have pointed to a crucial role of DOCK8 in the immune system, since the loss of DOCK8 function in lymphocytes leads to impaired immunity and an increased disposition for recurrent infections^29^ Particularly, DOCK8 regulates several functions of T cells, natural killer (NK) cells and B cells in humans and mice, by being essential for survival, immune synapse formation and activation of lymphocytes^18,20,30,31^. This critical involvement of DOCK8 in immune cell functions could in turn determine the role of septin 7 in the immune system, considering the fact that septins act as potential scaffolding platforms for interacting proteins, which mediate cellular functions.

Since the in-depth expression analysis of these interacting proteins is of considerable value for approaching cellular and molecular septin 7 functions, we characterized the expression pattern of DOCK8 in primary blood-derived lymphocytes (Figure 2). In controls, high levels of DOCK8 were detected in all lymphocytes, comparable to humans and mice^32^, pointing to a conserved role of DOCK8 in immune cells across species. However, we could detect a difference of DOCK8 protein levels between T cells and B cells in PBL of healthy controls. Whereas in lymphocytes of healthy humans, equal protein levels of DOCK8 were present in T cells and B cells^22^, our analysis revealed significant 2-fold higher (* = p ≤ 0.05) protein levels of DOCK8 in B cells compared to levels in CD4^+^ T cells (Figure 2).

Further, in our study DOCK8 levels significantly decreased in lymphocytes of autoimmune cases compared to healthy controls (Figure 3). Since DOCK8 is involved in several cellular processes in the immune system^18,20,30,33^, this significant decrease in lymphocytes of autoimmune cases might explain changes in the immune response in disease. So far, the effects of DOCK8 downregulation in a spontaneous autoimmune disease are not fully understood. But there were already some hints, that the clinical signs of DOCK8-deficiency can reach beyond typical symptoms of primary immune deficiencies. Particularly, in DOCK8-deficient individuals, a low prevalence of autoimmune manifestations was described, including vasculitis^29^, hemolytic anemia^34^ and even one case of uveitis suspected to be of autoimmune origin^35^. Hypotheses explaining the susceptibility of DOCK8-deficient patients towards autoimmunity included defective Treg function^36^. The selective DOCK8-deficiency in Tregs of FOXP3YFP-Cre/Dock8flox/flox mice revealed a higher risk to develop autoimmunity^37^, pointing to an important role of DOCK8 function in Tregs for maintaining self-tolerance. However, defective Treg functions in DOCK8-deficiency might occur due to secondary causes, since DOCK8-deficiency affects a wide range of immune cells^38^. Our results of DOCK8 decrease affecting different lymphocyte subsets in autoimmune cases, indicates that the presence of autoimmunity is not limited to selective impairment of DOCK8 functions in Tregs. In contrast, we suggest that the decreased DOCK8 expression in lymphocytes might result in autoimmunity due to a complex interplay of impaired T and B cell tolerance that correlates with respective findings in DOCK8-deficiency^36^.

Since our findings pointed to a crucial role of DOCK8 in autoimmune diseases, we were interested in the potential functional changes of lymphocytes in autoimmunity, driven by DOCK8. By detecting remarkable differences in DOCK8 protein-protein-interactions in lymphocytes of healthy and autoimmune cases, we demonstrated that DOCK8 takes a deviant role in disease. Since DOCK8 interaction partners were differentially enriched in both phenotypes and in general, more interacting proteins were identified from autoimmune PBL (178 interaction partners), we hypothesized that DOCK8 contributes to a deviant inter- and intracellular signaling in disease, resulting in changed immune response. Thus, we focused on differences in signaling pathways driven by DOCK8 and revealed changes in integrin signaling, which were associated with autoimmunity (Table 3). Proteins comprised in integrin pathway as talin 1 and integrin-linked kinase (ILK) (Table 3) determine crucial cellular responses as adhesion, migration or survival^25,39^ and we previously demonstrated their critical involvement in inflammatory processes of autoimmune cases in earlier studies^23,24,40^. Therefore, the enrichment of integrin pathway in lymphocytes of diseased specimen might explain autoreactive immune responses driven by DOCK8. Especially our finding of the DOCK8 interaction partner ILK, comprised in integrin pathway with enhanced enrichment in autoimmune cases (Table 3), reinforced this hypothesis, since this serine-threonine kinase regulates several immune functions^41,42^ ILK was clearly detected in DOCK8-immunoprecipitates and the reverse experiment showed the presence of DOCK8 in ILK-precipiates (Figure 4), verifying the protein-protein-interaction of both molecules. Interestingly, ILK was recently described to be specifically upregulated in PBL from horses with induced autoimmune uveitis during an acute uveitic attack^23^, which implicates a direct correlation of ILK upregulation and autoaggressive reaction of lymphocytes. Thus, we suggest that the increased ILK expression in lymphocytes of spontaneous autoimmune cases promotes changes in immune responses resulting in autoreactivity of activated lymphocytes. So far, cellular mechanisms driven by ILK overexpression in lymphocytes are not fully elucidated. However, a competitive advantage in trafficking and survival was demonstrated in murine ILK-competent T cells, whereas thymocytes of T cell-specific knockout mice showed diminished survival, particularly a higher apoptosis rate and increased necrosis after stressing of these cells^25^. Since ILK serves as an upstream modulator of Akt, which mediates inhibition of apoptosis^43^, the increased expression intensity of ILK in lymphocytes of autoimmune cases might result in higher survival, provoking insufficient deletion of autoreactive T-cells. Therefore, we determined apoptosis rate of primary blood-derived lymphocytes with spontaneously altered ILK expression (Figure 4). In lymphocytes of autoimmune cases, showing ILK overexpression, apoptosis rate decreased significantly compared to healthy specimen whereas necrosis was consistently negligible in both phenotypes (< 1%) (Figure 4), indicating a correlation between changed ILK expression and reduced apoptosis in ERU. These interesting findings will be profoundly analyzed in subsequent studies to elucidate the impact of divergent apoptotic behavior of cells on autoreactive immune responses in disease.

In summary, our findings provide deeper insight into interaction networks in lymphocytes and contribute to a better understanding of immune cell functions. Moreover, we gained information about the role of septin 7 and its novel interactor DOCK8 in autoimmune disease. We propose that the decreased expression levels of these potential key players in autoimmune uveitis and the deviant signaling pathways driven by DOCK8 promote autoreactive inflammatory processes.

## METHODS

### Sample preparation

For this study, PBL of 20 healthy and 54 ERU cases were used. Autoimmune cases included in this study had suffered from at least three uveitic attacks and were diagnosed by clinical signs of acute uveitis accompanied by a documented history of recurrent intraocular inflammation^44^ No experimental animals were used in this study. Horses were treated according to the ethical principles and guidelines for scientific experiments on animals according to the ARVO statement for the use of animals in Ophthalmic and Vision research https://www.arvo.org. The study protocol to obtain blood samples from ERU cases in quiescent stage of disease and controls (both obtained from Equine Clinic, LMU Munich, Germany) was permitted by the Ethical Committee of the local authority (Regierung von Oberbayern; permit number: ROB-55.2Vet-2532.Vet_03-17-88). All experiments were performed in accordance with the relevant guidelines and regulations.

In detail, mass spectrometric analysis of septin 7 interaction network was performed in PBL of a healthy control. Additionally, PBL of eight healthy horses were used for confirming septin 7 interaction partners by co-immunoprecipitation. Phenotyping DOCK8 expression in healthy controls and ERU horses was achieved by flow cytometric analysis of 13 controls and 12 ERU cases and by western blot analysis of 16 healthy and 36 ERU horses. PBL of three healthy and three ERU cases were used for co-immunoprecipitation of DOCK8 followed by mass spectrometric analysis. Expression analysis of ILK by western blot was performed in 17 controls and 34 ERU cases. Apoptosis and necrosis rates were investigated in four healthy and six ERU horses.

Equine venous blood samples were collected in tubes filled with heparin sodium 25.000 I.U. (Ratiopharm, Ulm, Germany) diluted in RPMI (PanBiotech, Aidenbach, Germany). PBL from plasma were isolated by density gradient centrifugation (room temperature (RT), 290 × g, 25 min, brake off) using Pancoll separation solution (PanBiotech). Lymphocytes were extracted from intermediate phase. Cells were either used immediately or stored at -20°C (lysate) or -80°C (vital cells).

### Immunoprecipitation (IP) of interacting proteins

5 × 10^8^ PBL derived from whole blood were lysed in IP-lysis-buffer (TBS containing 50 mM Tris and 150 mM NaCl, 2% CHAPS, Roche complete EDTA-free protease inhibitor) and protein concentration was determined by use of Bradford protein assay (Sigma-Aldrich, Taufkirchen, Germany). 40 μl of Protein G Sepharose Beads (GE Healthcare, Freiburg, Germany) per 1 mg protein were washed with IP-buffer diluted with TBS and incubated separately with rat anti-septin 7 antibody (clone 19A4, self-made monoclonal rat anti-septin 7 antibody, isotype IgG2c; neat), rabbit polyclonal anti-DOCK8 antibody (H-159, Santa Cruz, Heidelberg, Germany; 6 μg/ml) diluted in IP-buffer or respective isotype controls were used. Lysates were added to beads and incubated on a tube rotator over night at 4°C. Beads were washed with IP-lysis-buffer before immunoprecipitates were eluted by adding Laemmli-buffer.

### Sample preparation for LC-MS/MS mass spectrometry

Laemmli eluates were diluted in TBS 1:4 and 10 μl 100 mM dithiothreitol was added for 30 min at 60°C. After cooling down, 100 μl UA-buffer (8 M urea + 1 M Tris-HCl pH 8.5 diluted in HPLC-grade water) was added. Eluates were transferred to 30 kDa cut-off centrifuge filters (Sartorius, Göttingen, Germany) and washed with 200 μl UA-buffer and with 100 μl ABC-buffer (50 mM NH_3_HCO_3_ diluted in HPLC-grade water). After washing, the proteins were subjected to proteolysis at RT for 2 h with 1 μg Lys-C in 50 μl ABC-buffer followed by addition of 1 μg trypsin in 10 μl of ABC-buffer and incubation at 37°C overnight. Peptides were collected by centrifugation and acidified with 0.5% trifluoroacetic acid.

### Mass spectrometry for identification of interacting proteins

LC-MS/MS analyses were performed on an LTQ OrbitrapXL (Thermo Fisher Scientific, Bremen, Germany). Briefly, peptides were loaded onto an Ultimate3000 nano HPLC system (Dionex, Idstein, Germany) equipped with a nano trap column at a flow rate of 30 μl/min in HPLC-buffer. Peptides were separated on a reversed phase chromatography over 140 min at a flow rate of 300 nl/min. For mass spectrometric analysis, a maximal injection time of 100 ms for MS spectra and 500 ms for MS/MS spectra was defined. MS spectra were recorded within a mass range from 200-2000 Da at a resolution of 60,000 full-width half-maximum. All ions which were selected for fragmentation (top 10 method) went through dynamic exclusion over 30 sec.

### Protein identification and label-free quantification

Acquired MS spectra were imported into Progenesis software (version 2.5 Nonlinear Dynamics, Waters and analyzed as previously described^45,46^. After alignment, peak picking, exclusion of features with charge of 1 and >7 and normalization, spectra were exported as Mascot Generic files and searched against the Ensembl Horse database (Version 78) with Mascot (Matrix Science, Version 2.4.1). Peptide assignment was reimported to Progenesis Software. All unique peptides allocated to a protein were considered for quantification. Only proteins quantified with at least two peptides were included for further analysis. Abundances in individual samples were calculated by adding the single abundances of the respective peptides. Protein ratios between septin 7- or DOCK8-IP samples and the respective isotype control samples were determined by dividing these abundances. Data of DOCK8 interaction proteomics was analyzed with gene ranker (Genomatix version 3.7) for enrichment of canonical signal transduction pathways, using orthologous genes.

### Western blots

1 × 10^7^ PBL were lysed in lysis buffer (9 M Urea, 2 M Thiourea, 1% Dithioerythritol, 4% CHAPS, 2.5 μM EDTA) and Laemmli-buffer was added. Co-IP-eluates were just mixed with Laemmli-buffer before gel-electrophoresis. Protein expression was analyzed separately for every biological replicate of controls (n=16) and autoimmune cases (n=36). From each sample, 7 μg protein was separated by SDS-PAGE on 8-10% gels and blotted semidry onto PVDF membranes (GE Healthcare). To prevent unspecific binding, membranes were blocked with 4% BSA or 1% PVP-T. After washing, blots were incubated with rat anti-septin 7 monoclonal antibody (clone 19A4, self-made; neat), anti-DOCK8 (rabbit polyclonal, Santa Cruz Biotechnology; 1:1000), anti-ILK (mouse monoclonal, Santa Cruz Biotechnology; 1:1000) and anti-beta actin (mouse, monoclonal, Sigma-Aldrich; 1:50000) to control equal loading at 4°C overnight. Secondary antibodies were taken respectively. HRP-conjugated anti-rabbit IgG antibody (Cell Signaling Technology, Darmstadt, Germany; 1:10000), HRP-conjugated antirat IgG 2c antibody (1:8000) or HRP-conjugated anti-mouse IgG antibody (Sigma-Aldrich; 1:5000) were used for incubation at RT for one hour. After six washing steps, signaling was detected by enhanced chemoluminescence on X-ray film (SUPER-2000G ortho, Fuji; Christiansen, Planegg, Germany). Quantification of western blot signals was achieved by using open source Image J software https://imagej.net/. All abundances were normalized to respective beta actin signals.

### Analysis of DOCK8 expression by flow cytometry

Staining of 5 × 10^5^ cells per well was performed with anti-equine CD4 (mouse IgG1 monoclonal, Bio-Rad AbD Serotec, Puchheim, Germany; 1:100), PE-conjugated anti-equine CD8 (mouse IgG 2a monoclonal, Bio-Rad AbD Serotec; 1:10) and anti-equine B cells (clone CVS 41, monoclonal mouse IgG anti-horse Ig light chain, Bio-Rad AbD Serotec; 1:10) antibodies, diluted in FACS staining buffer (1% BSA + 0,001% NaN_3_ in PBS). Staining with anti-DOCK8 antibody (rabbit polyclonal, Santa Cruz; 1:10) was performed after permeabilization of cells (BD Cytofix/Cytoperm fixation/permeabilization kit; BD Biosciences, Heidelberg, Germany). For all stainings, respective isotype controls were used. Respective secondary antibodies anti-mouse IgG1 PE, anti-mouse IgG PE (both Santa Cruz Biotechnology; 1:300) and anti-rabbit IgG Alexa 488 antibody (Invitrogen, Karlsruhe, Germany; 1:400) were added. Cells were fixed in 1% PFA diluted in staining buffer and stored at 4°C until further processing. On FACS Canto II, measurement of cells was performed with FACS Diva Software (BD Biosciences). Lymphocytes were gated according to forward scatter (cell size) and side scatter (intercellular granularity) properties of cells. Per staining, between 5 × 10^3^ to 1 × 10^4^ cells were measured per staining. Further analysis of flow cytometry data was performed using open source Flowing Software 2.5.1 http://flowingsoftware.btk.fi/ (Perttu Terho, Turku Centre for Biotechnology, Finland).

### Analysis of cell apoptosis

To investigate the effects of ILK expression differences on apoptosis, 1x10^7^ PBL from healthy and ERU cases were cultured in RPMI containing 6% fetal bovine serum (both PAN-Biotech). To compare apoptosis and necrosis between PBL of healthy controls and ERU horses, the Annexin V FITC Apoptosis Detection Kit (Sigma-Aldrich) was used on 1–10^5^ cells in a 96-well-plate. Cells were incubated with 2 μl Annexin V-FITC and 8 μl propium iodide (PI) for 10 minutes. The amount of apoptotic (annexin V positive) und necrotic cells (PI positive) was measured immediately after incubation time by using flow cytometry.

### Statistics

For determination of Gaussian distribution, Kolmogorow-Smirnow (KS) test was used. If KS test was significant (p ≤ 0.05; normal distribution), student’s *t*-test was used for statistical analysis, if KS test was not significant (p > 0.05; no normal distribution) statistics were performed using Mann-Whitney test. Statistical comparisons of DOCK8 expression intensities in lymphocyte subsets were performed by using student’s *t*-test. For comparison of DOCK8 expression intensity in healthy controls, DOCK8 geo MFI values were presented as means ± SD, for comparing DOCK8 expression in healthy and diseased specimen, controls were set to a 100% ± SD. Statistical analyses of DOCK8 expression intensity differences and ILK expression intensity differences in western blots as well as the comparison of apoptosis and necrosis rates were performed by using Mann-Whitney-test. In both tests, statistical probabilities were considered significant at p ≤ 0.05.

### Data Availability

The datasets generated during and/or analyzed during the current study are available from the corresponding author on reasonable request.

## ACKNOWLEDGMENTS

This work was supported by a grant from the Deutsche Forschungsgemeinschaft DFG DE 719/4-3 (to C.D.). The authors would like to thank Hartmut Gerhards and Lutz S. Göhring as well as the staff of the Equine Clinic at LMU Munich for providing blood samples.

## AUTHOR CONTRIBUTIONS

C.D. conceived, designed, performed and analyzed the experiments and supervised the project; M.S., S.H., C.W. and K.K. performed the experiments; M.S., S.H., C.W., R.D. and K.K. analyzed the data; S.H. contributed reagents, materials and analysis tools; E.K. produced antibodies; M.S. and C.D. wrote the manuscript. All authors critically read the manuscript and approved the final version to be published.

